# Mitochondrial NAD-malic enzyme subunits modulate early fungal penetration and differentially shape the *Arabidopsis* proteome

**DOI:** 10.64898/2026.11.20.768030

**Authors:** Maria del Pilar Martinez, Alexander Fray, Ioana Nica, Noah Ditz, Julia Seufer, Christiane Rohrbach, Philipp Westhoff, Holger Eubel, Lars M. Voll, Veronica G. Maurino

## Abstract

Mitochondrial NAD-dependent malic enzyme (NAD-ME) couples organic acid metabolism to cellular redox balance through the conversion of malate to pyruvate, accompanied by NADH production. In *Arabidopsis thaliana*, NAD-ME is mainly a heterodimer of two distinct subunits, ME1 and ME2, which their individual physiological roles remain poorly understood. In this study, we use *me1*, *me2* and *me1/me2* mutant lines to investigated the contributions of both subunits to metabolic homeostasis, proteome organization, and susceptibility to the hemibiotrophic fungus *Colletotrichum higginsianum*. Under standard growth conditions, loss of ME2 reduces ME1 protein abundance to approximately 20% of wild-type levels despite sustained ME1 transcript accumulation, whereas ME2 protein abundance remained at approximately 75% of wild-type levels in the absence of ME1. These findings suggest an asymmetric interdependence between NAD-ME subunits, with ME2 playing a role in maintaining ME1 protein abundance. Consistent with these differences, the basal proteome of *me1* plants closely resembled that of wild type, whereas *me2* and *me1/me2* exhibit extensive and overlapping proteomic changes, particularly reduced abundances of photosynthesis-associated proteins, without detectable impairment of photosynthetic performance. Diel metabolite profiling revealed selective changes in organic acid and sugar levels, indicating distinct contributions of the ME subunits to primary metabolism. Interestingly, loss of either one, or both subunits, significantly reduces penetration rates of the fungal pathogen *C. higginsianum*. Despite this shared interaction phenotype, fungal infection induces extensive proteomic remodeling in *me1*, whereas comparatively few changes occurred in *me2* and *me1/me2*. Notably, several proteins reduce upon infection in *me1* are already less abundant in non-infected ME2-deficient plants. Together, these findings highlight the distinct contributions of NAD-ME subunits to proteome organization and metabolic homeostasis and suggest that NAD-ME influences cellular conditions that favor successful fungal penetration.

## INTRODUCTION

Mitochondrial NAD-dependent malic enzyme (NAD-ME; EC 1.1.1.39) catalyzes the oxidative decarboxylation of malate to pyruvate and CO_2_, coupled to NAD^+^ reduction. Its activity links malate turnover to the production of reducing-equivalent in the mitochondrial matrix and may influence the allocation of carbon and reducing equivalents between cellular compartments. NAD-ME derived pyruvate was initially considered a major substrate pool for the mitochondrial pyruvate dehydrogenase complex (PDC), but recent studies indicate that pyruvate imported through the mitochondrial pyruvate carrier and pyruvate produced by NAD-ME can enter distinct metabolic pathways (Le et al., 2021; Le et al., 2022). Imported pyruvate is preferentially oxidized by PDC under the conditions examined, whereas NAD-ME-derived pyruvate is preferentially exported or trans-aminated to alanine. Such compartmentation suggests that NAD-ME influences coordination of pyruvate metabolism beyond respiration, although the contribution of its pyruvate product to downstream cytosolic or plastid pathways remains unresolved. In addition, NAD-ME-derived NADH may contribute to intercompartmental transfer of reducing equivalents through metabolite-based shuttles, such as the malate-oxaloacetate shuttle, thereby potentially linking mitochondrial malate metabolism to cellular redox homeostasis (Selinski and Scheibe, 2019).

In *Arabidopsis thaliana*, mitochondrial NAD-dependent malic enzyme (NAD-ME) is encoded by two genes, *ME1* (*At2g13560*) and *ME2* (*At4g00570*). Their protein products, ME1 and ME2, can assemble into enzymatically active homo– and heterodimers, with the heterodimer being the predominant form *in vivo* (Tronconi et al., 2008; Tronconi et al., 2020). Biochemical characterization of recombinant NAD-ME proteins revealed distinct properties depending on their subunit composition (Tronconi et al., 2010). The recombinant ME1 homomer shows sigmoidal malate kinetics and activation by fumarate, whereas the ME2 homomer displays a hyperbolic response to malate and is activated by CoA, phosphoenolpyruvate, and fructose 1,6-bisphosphate. The ME1-ME2 heterodimer responds synergistically to fumarate and CoA. These findings support biochemical specialization but do not establish whether active homodimers operate *in vivo*. Furthermore, ME1 and ME2 also differ in their contributions to total NAD-ME activity and tissue distribution, with ME2, but not ME1, detected in anthers (Tronconi et al., 2008; Tronconi et al., 2010). Mutants lacking either or both ME subunits retain normal vegetative growth and photosynthetic capacity under favorable conditions (Tronconi et al., 2008; Martinez et al., 2026). However, the absence of an obvious phenotype does not rule out distinct contributions of the two subunits to metabolism and stress responses. Interestingly, *me1* mutants retain approximately 50% of wild-type activity and *me2* mutants less than 20% (Tronconi et al., 2008; Tronconi et al., 2010), suggesting that ME2 can sustain NAD-ME-associated functions without ME1 more effectively than in the reverse situation. However, the significance of these differences between ME1 and ME2 remains largely unexplored.

Alterations in mitochondrial respiratory metabolism can influence plant immune responses by modulating cellular redox homeostasis and reactive oxygen species (ROS) production. Recent evidence has linked mitochondrial NAD-ME to plant–pathogen interactions. Silencing NAD-ME1 in *Nicotiana benthamiana* promoted Bamboo mosaic virus accumulation, altered cytosolic NADH/NAD⁺ balance, and affected defense-related gene expression (Hou et al., 2025). More recently, Barreto et al. (2026) demonstrated that mitochondrial NAD-ME contributes to early metabolic and redox responses during pattern-triggered immunity in *A. thaliana*. Loss of both NAD-ME subunits attenuated the flg22-induced reduction of the cellular NAD pool and suppressed the intracellular glutathione redox response, while leaving the extracellular ROS burst unaffected. These findings implicate NAD-ME in the metabolic reprogramming associated with early immune signaling. However, the distinct contributions of the two mitochondrial NAD-ME subunits to plant interactions with fungal pathogens remain largely unexplored.

We therefore investigated how the individual and combined loss of ME1 and ME2 affects mitochondrial NAD-ME subunit abundance, proteome composition and metabolic organization in *A.s thaliana*, as well as susceptibility to the hemibiotrophic fungus *Colletotrichum higginsianum*. To address these questions, we analyzed transcript and protein abundance, basal proteomic and metabolic profiles, photosynthetic performance, fungal penetration, and infection-associated proteomic responses. By comparing single and double mutants, we aimed to identify both shared and subunit-specific contributions to cellular metabolism, proteome organization, and plant-pathogen interactions.

## RESULTS

### *ME1* and *ME2* show distinct diel transcript profiles

To elucidate the transcriptional interplay between *ME1* and *ME2*, we first examined whether the loss of one NAD-ME subunit affects the expression of the other. Transcript abundance was quantified by RT-qPCR in the *me-*single mutants, *me1* and *me2.1* (*me2*), which we previously characterized as knockout lines (Suppl. Table 1) (Tronconi et al., 2008; Martinez et al., 2026). To assess diurnal changes in gene expression, whole rosettes were harvested at three time points during the photoperiod: one hour before lights-on (−1), one hour after lights-on (+1), and one hour before lights-off (+7).

In the *me2* line, *ME1* transcripts are increased relative to wild type (Col-0) one hour after dawn (+1) but not at the end of the night (−1) or near dusk (+7) with respect to wild type (Fig. 1A). Conversely, ME2 transcripts in *me1* are increased at the end of the night (−1), but reduced during the light period (+1 and +7) with respect to wild type (Fig. 1A). Thus, loss of either subunit affects the other’s expression in a time-dependent manner, effect on *ME1* is strongest in *me2* at dawn, while effect on *ME2* is strongest in *me1* around dawn, rather than producing a uniformly compensatory transcriptional response.

**Figure 1.**
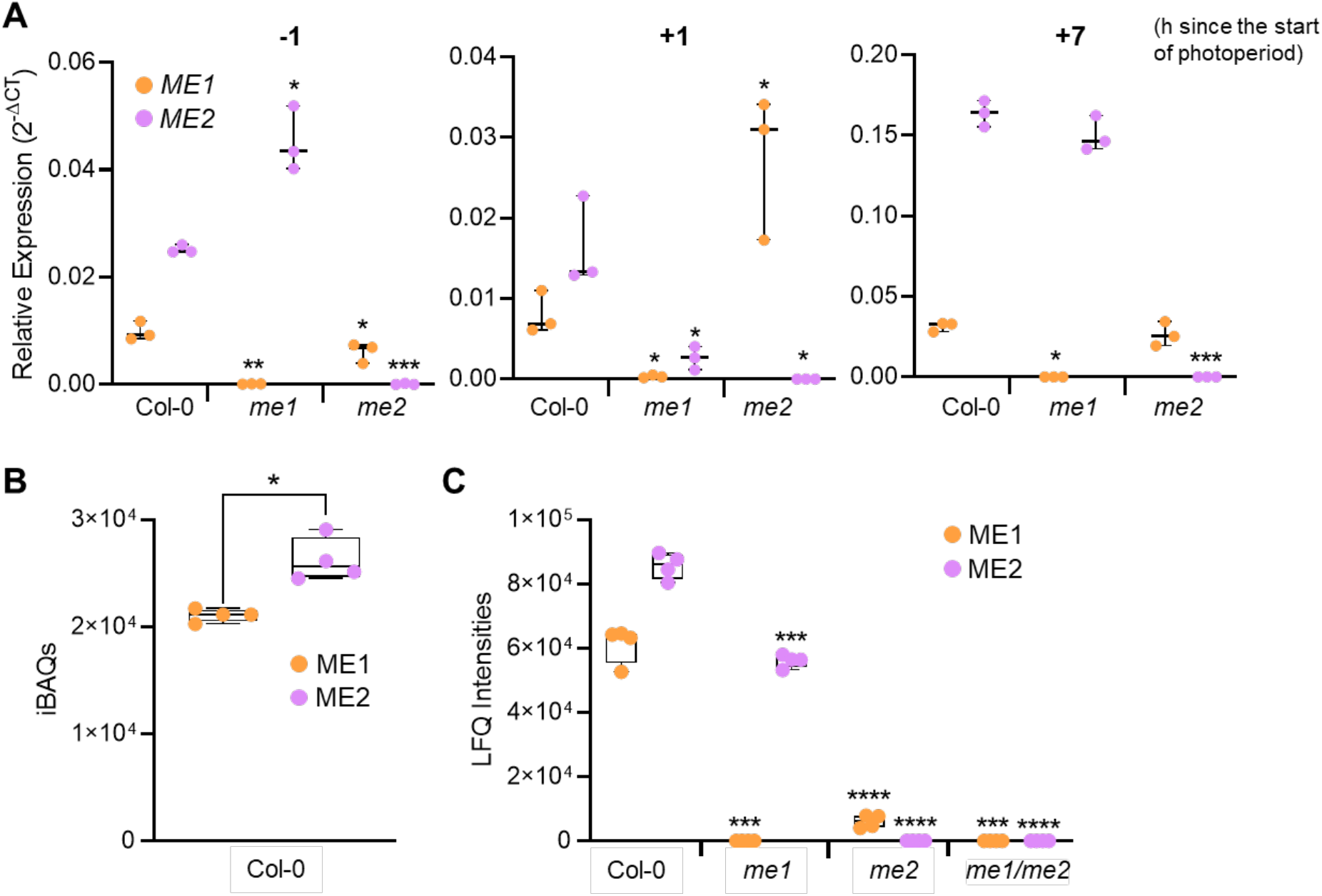
*A. thaliana ME* transcript and protein abundances in *me*-mutant lines and Col-0. **A)** Transcript levels of *ME1* and *ME2* were analyzed in the different mutants. Samples were collected one hour before the onset light period (−1), one hour after the onset of the light period (+1), and one hour before the end of the light period (+7). Statistical significance between the mutants and Col-0 was determined using Welch’s t-test (p < 0.05, *; p < 0.01, **; p < 0.001, ***; p < 0.0001, ****). **B)** Intensity-Based Absolute Quantification (iBAQs) values of ME1 and ME2 in Col-0 and **(C)** Label-Free Quantification (LFQ) intensities of ME1 and ME2 in Col-0 and mutant lines grown under LL-SD (control condition). Statics between the abundances of ME1 and ME2 in (A) and between Col-0 and the *me*-mutant lines in (B) was assessed by Welch’s t-test (p < 0.05, *; p < 0.01, **; p < 0.001, ***; p < 0.0001, ****).

In wild-type plants, *ME2* transcripts are more abundant than *ME1* transcripts at the sampling times examined (−1, +1 and +7), and both transcripts are more abundant towards the end of the light period (−1). The difference is greatest near the end of the light period (+7), when *ME2* transcript abundance exceeds *ME1* by more than threefold (Fig. 1A), indicating distinct diel expression patterns.

To investigate potential differences in transcriptional regulation, we used PlantCARE to identify putative cis-regulatory elements within the 2-kb genomic regions upstream of the *ME1* and *ME2* translation start codons (Lescot et al., 2002). Both promoters contain predicted motifs associated with light, ABA, and stress responses, including ABRE, G-box, MYB/MYC, STRE, W-box, and as-1 elements (Suppl. Fig. 1; Suppl. Table 3). Several predicted motifs differ between the promoters: a DRE core, TCT motif, and Box S motifs were identified only in ME1, whereas AP-1 and HD-Zip-1 motifs were identified only in ME2. These differences in predicted cis-regulatory elements suggest that *ME1* and *ME2* may be subject to distinct transcriptional regulatory mechanisms, potentially contributing to their differential expression patterns across the diurnal cycle.

### ME2 is required for normal ME1 protein accumulation

To assess whether changes in transcript abundance are reflected at the protein level, we semi-quantified ME1 and ME2 abundances by shotgun mass spectrometry in the *me1* and *me2* single mutants, and the *me1/me2* double mutant. As no major changes in protein abundance are expected over the diurnal cycle (Lee et al., 2010), we therefore selected a single harvest time point to assess protein abundance: one hour before the end of the photoperiod (+7).

DIA-PASEF (data-independent acquisition with parallel accumulation-serial fragmentation) was employed as the mass spectrometry acquisition method (Meier et al., 2020). This approach facilitates accurate semi-quantitative abundance estimations, with 17-19 ME1-unique peptides, and 21-22 ME2-unique peptides identified across samples, enabling reliable quantification of both NAD-ME proteins. In wild-type plants, ME2 is more abundant than ME1 (Fig. 1B). In *me1*, ME2 abundance is reduced to approximately 75% of wild-type level, whereas in *me2*, ME1 abundance decreases markedly to approximately 20% of wild type (Fig. 1C). Interestingly, this pronounced reduction in ME1 protein abundance contrasted with the increased accumulation of *ME1* transcripts observed in *me2* at the beginning of the light period.

These results establish a strongly asymmetric dependence of subunit protein accumulation: most ME2 remains detectable without ME1, whereas ME1 abundance is markedly reduced without ME2. Because *ME1* transcripts remain present in *me2*, the effect is likely to involve post-transcriptional control. Together with prior enzymatic activity measurements (Tronconi et al., 2008), the persistence of ME2 in *me1* is compatible with residual ME2-dependent NAD-ME activity.

### Loss of ME2 causes extensive cellular proteome remodeling resembling the situation in the double mutant

We next compared protein abundances in wild type and the three mutant genotypes, using samples collected on hour before the end of the day (+7) (Fig. 2). Principal component analysis (PCA) grouped *me1* with wild type, whereas *me2* clustered with *me1/me2* (Fig. 2A). Relative to wild type, no protein groups differed significantly in *me1*. In contrast, 318 groups differed between *me2* and wild type (157 higher and 161 lower), and 264 differed in *me1/me2* (140 higher and 124 lower) (Fig. 2B). The marked similarity of *me2* and *me1/me2* indicates that ME2-loss has a larger impact on basal proteome organization than ME1-loss, possibly because in *me2* ME1 abundance is very low mimicking a double mutant. Overlap analysis further supported this distinction. The *me2* and *me1/me2* mutants share 48 lower-abundance and 56 higher-abundance protein groups relative to wild type (Fig. 2C).

**Figure 2.**
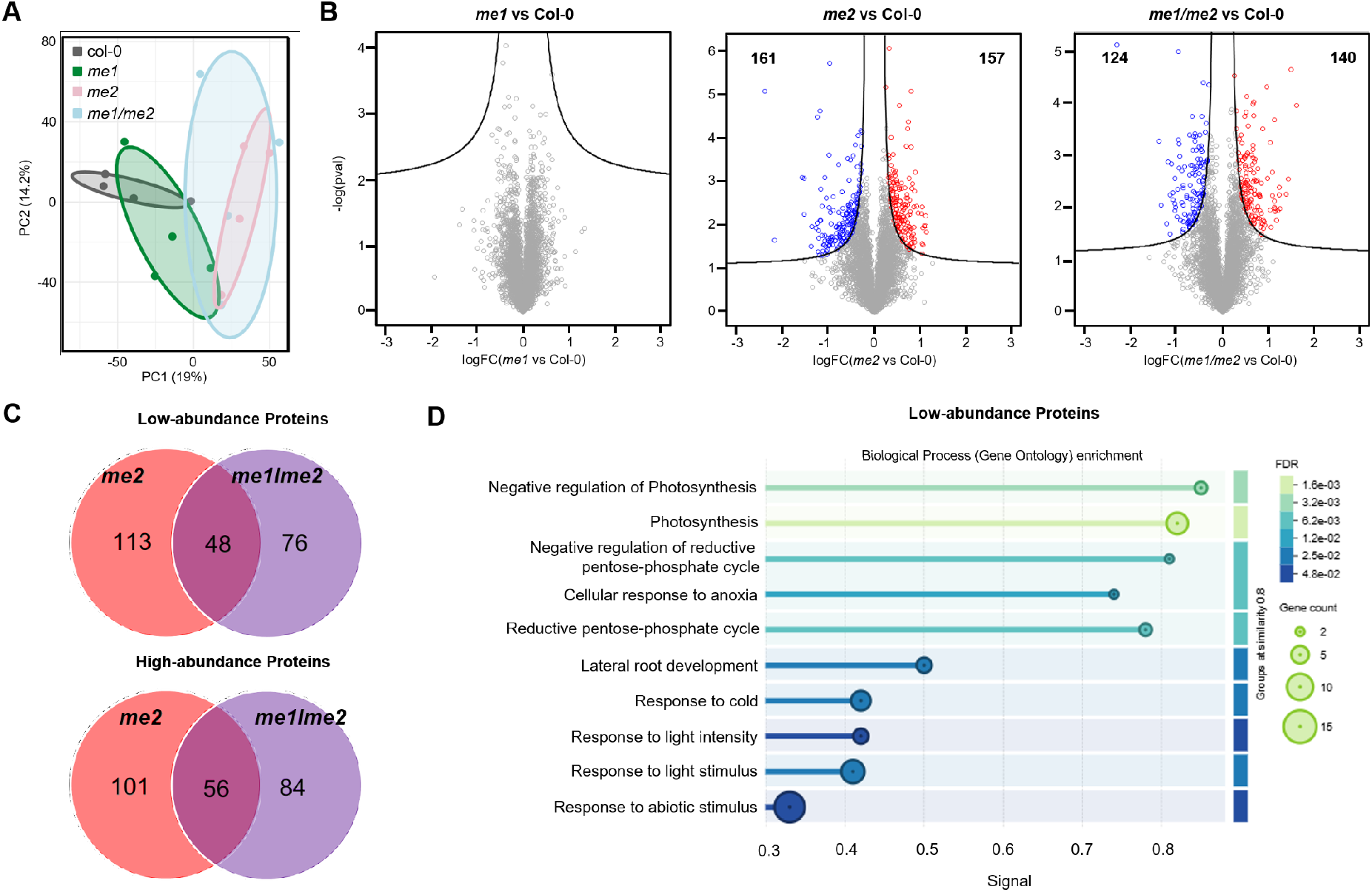
Proteomic analysis of *me* mutant lines and Col-0. **A)** PCA analysis of *me1*, *me2*, *me1/me2* and Col-0 grown under SD-LL (n=4). **B)** After normalization, log_2_-transformed LFQ values of protein groups were used to assess differences in the whole proteome between *me1*, *me2* and *me1/me2* against Col-0 by statistical testing employing four independent biological replicates. The complete list of categories is provided in Supplemental Data 1. Protein groups located in the areas above the black lines fulfil the significance threshold limits (FDR ≤ 0.05; S0 = 0.1). Red dots represent protein groups with higher abundance in the mutant line whereas blue indicates lower abundance protein groups. **C)** Unique and common protein groups with higher and lower abundance in the mutant lines against Col-0. **D)** Gene Ontology enrichment using STRING (v12.5) with common low abundance proteins between *me2* and *me1/me2* selected in **C**.

Gene Ontology (GO) analysis using STRING (v12.5; Szklarczyk et al., 2025) of the 48 proteins decreased in both *me2* and *me1/me2* relative to the wild type highlighted the light reaction of photosynthesis and the Calvin-Benson cycle (Fig. 2D). Rubisco activase (RCA) is down to approximately 20 % of wild type levels in these mutants; CP12-1 and CP12-2 are also lower abundant in *me2* and *me1/me2*, presenting approximately 30% and 50 % of wild type levels, respectively. Additional decreases affected components of photosystems I and II. The common set of proteins with higher-abundance in *me2* and *me1/me2* compered to Col-0 was not significantly enriched in any biological process by GO. These results indicate that photosynthesis-associated proteins are particularly affected in the *me2* single and *me1 me2* double mutants, both of which exhibit strongly reduced abundance of both NAD-ME subunits.

### Photosynthetic performance is maintained in *me2* and *me1/me2* mutants despite changes in protein abundance

Because several proteins involved in photosynthesis are less abundant in *me2* and *me1/me2*, the photosynthetic performance of wild type and the three mutant lines was assessed. We found that maximum PSII quantum efficiency (Fv/Fm), PSII quantum yield (ΦPSII), electron transport rate (ETR), and non-photochemical quenching (NPQ) do not differ significantly among genotypes (Suppl. Fig. 2). Thus, the reduction in abundance of selected photosynthetic proteins did not translate into detectable impairment in the measured parameters under these conditions, where also no obvious differences in plant growth were observed relative to wild type (Martinez et al., 2026).

### Diel metabolite profiles reveal selective effects on organic acid and sugar pools

To examine whether the subunit-dependent proteome changes are accompanied by altered metabolism, we profiled soluble metabolites by GC-MS in whole rosettes. Samples were collected one hour before dawn (−1), one hour after dawn (+1), and one hour before dusk (+7). PCA did not reveal pronounced genotype-specific separation at these time points (Suppl. Fig. 3A), suggesting that the overall metabolite profile is comparatively stable despite the loss of one or both NAD-ME subunits.

Nevertheless, several metabolites differ among the lines. At the end of the night, malate is more abundant in *me2* than in *me1* (Fig. 3A). At the same time point, citrate and isocitrate are higher in *me1* than in *me1/me2*, with intermediate values in *me2*. Succinate is lower in *me1* than in the other genotypes one hour before and after dawn, while *me2* and *me1/me2* accumulate more succinate than wild type at the end of the night (Fig. 3A). Sugar changes are modest. At +1, hexoses are higher in *me2* and *me1/me2* than in *me1* and wild type. By +7, their abundance in *me2* approaches wild-type values, whereas the double mutant retains increased levels. Sucrose levels do not differ significantly among genotypes at the examined times (Fig. 3B).

**Figure 3.**
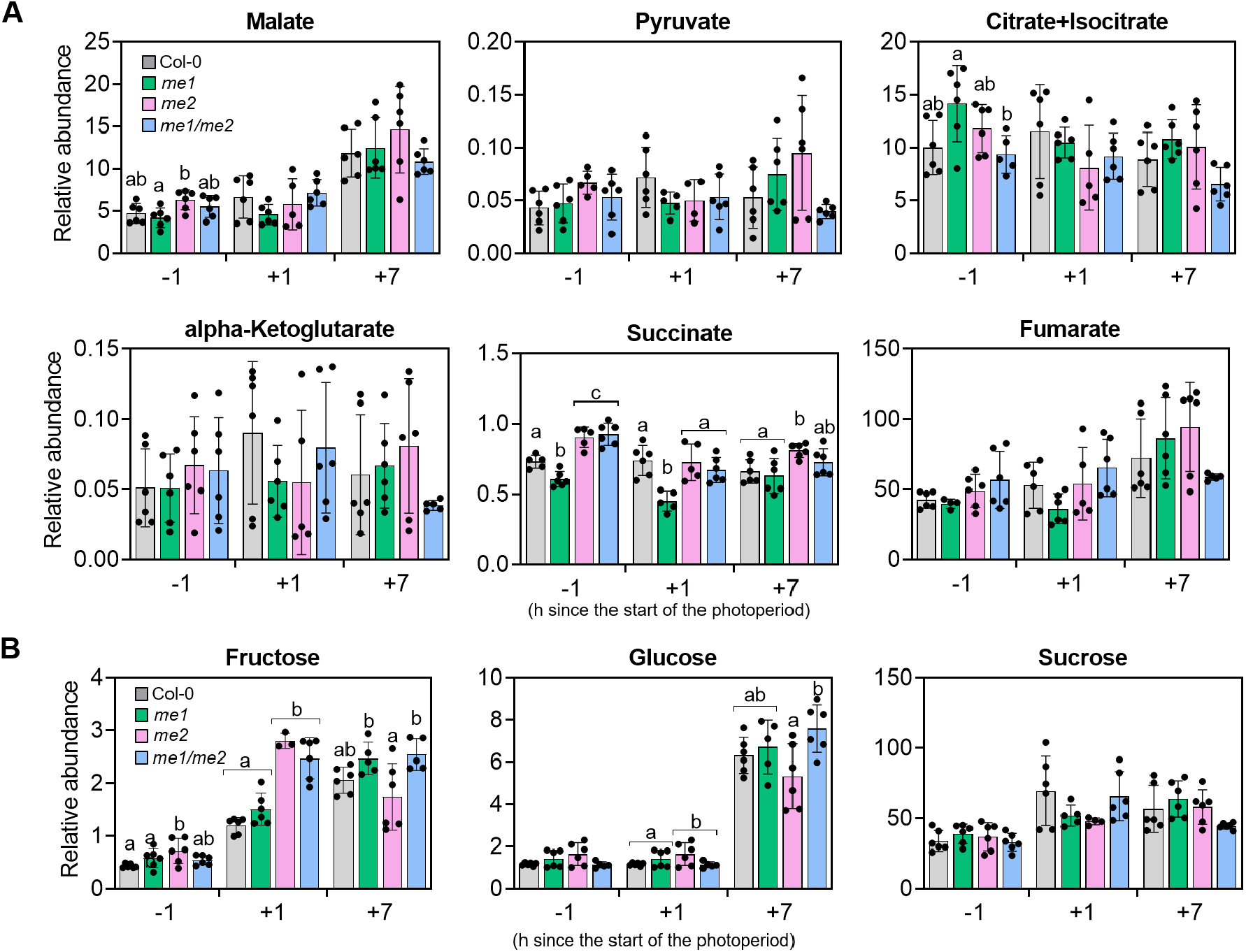
Relative metabolite abundances in leaves of plants grown in SD at LL. Relative levels of **(A)** TCA cycle intermediates and **(B)** sugars. Samples were harvested 1 h before lights-on (−1 h), 1 h after lights-on (+1 h) and 1 h before lights-off (+7 h). Bars show means ± SE (n = 6) independent biological replicates). Statistics were performed for each time point using one-way ANOVA followed by Tukey’s post-hoc test; bars that do not share a letter differ significantly at *p* < 0.05. No letter indicates no statistically significant differences between genotypes at the given time point.

Overall, these results indicate that loss of NAD-ME subunits causes selective, time-dependent changes in organic acid and sugar metabolism, with more pronounced alterations in *me2* and *me1/me2*, despite relatively modest effects on the overall metabolite profile.

### Disruption of either NAD-ME subunit reduces fungal penetration success

Previous work revealed that impairing mitochondrial metabolism by compromising complex I and II affected the interaction with the hemibiotrophic fungus *C. higginsianum* (McCollum et al., 2019). Given the distinct proteomic and metabolic alterations observed in the NAD-ME mutants, particularly changes in succinate abundance, we next investigated whether loss of NAD-ME subunits also influences susceptibility to fungal infection. To this end, we challenged plants with *C. higginsianum*. Fungal structures were quantified at 2.5 days post-inoculation, distinguishing appressoria from biotrophic and necrotrophic hyphae to assess fungal penetration success (Fig. 4).

**Figure 4.**
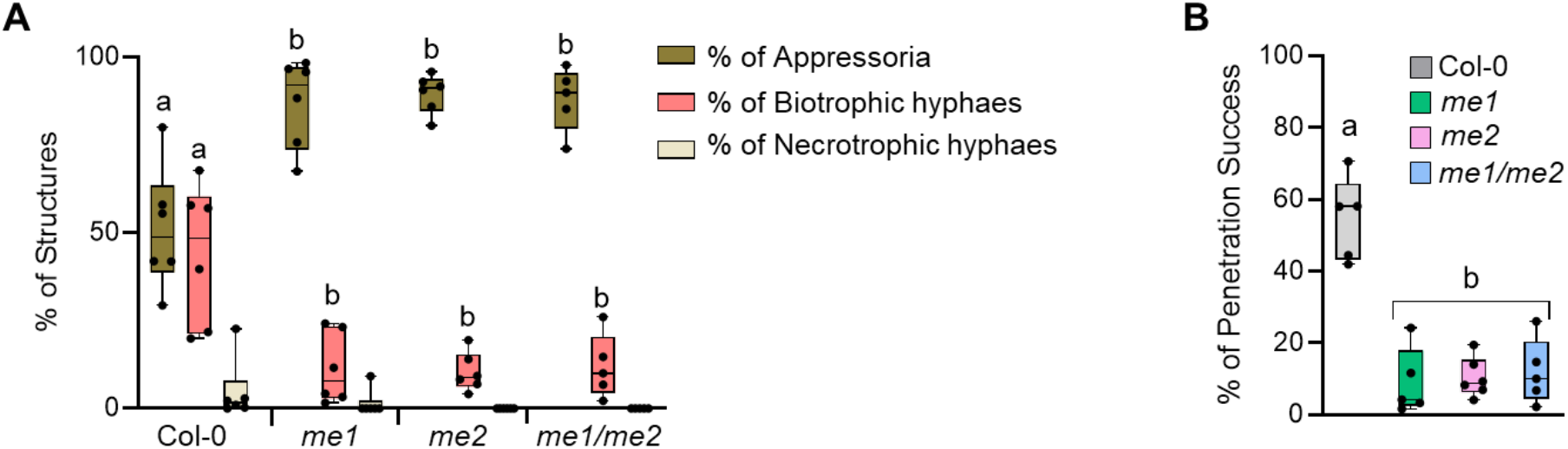
Susceptibility of Col-0 and mutant lines against *C. higginsianum* during early infection stage. **A)** Fungal structures were quantified 2.5 days after inoculation. Interleaved box and whiskers showing minimum to maximum values (n = 6, with approx. 100 penetration sites analyzed per replicate). **B)** Percentage of penetration success at 2.5 days after infection. Statistics were performed using one-way ANOVA followed by Tukey’s post-hoc test. Different letters indicate statistically significant differences (p < 0.05).

At 2.5 days post-inoculation, *C. higginsianum* on *me1*, *me2*, and *me1/me2* leaves is still in the appressorial stage to a larger extent and developed fewer biotrophic hyphae than on wild type leaves (Fig. 4A and B). The proportion of fungal structures associated with successful penetration is therefore significantly lower in single and double mutants. This shared phenotype indicates that perturbation of either NAD-ME subunit reduces early susceptibility to *C. higginsianum*.

### The proteomic response to infection differs between ME1– and ME2-deficient plants

To investigate whether the loss of mitochondrial NAD-ME subunits affects early host responses to fungal infection, we compared the proteomes of infected and mock-treated plants at two days post-inoculation. The previously observed differences in NAD-ME subunit abundance are maintained following infection. ME2 remains detectable in *me1*, whereas ME1 abundance is strongly reduced in *me2*, independently of treatment (Fig. 5B). In wild-type plants, ME1 is less abundant than ME2 under mock conditions, but their abundances became similar following infection (Fig. 5A). This suggests that infection may influence the relative abundance of NAD-ME subunits.

**Figure 5.**
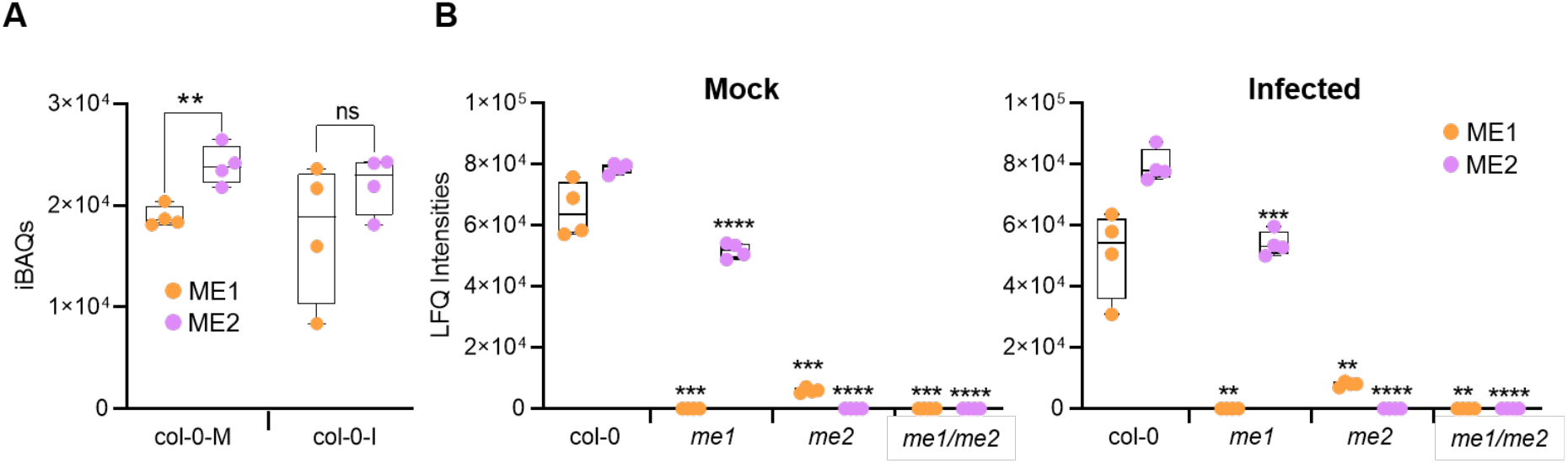
ME1 and ME2 abundances under mock and infected leaves of Col-0 and *me* mutant lines. **(A)** iBAQs values of ME1 and ME2 in infected (I) and mock (M) Col-0 and **(B)** Label-Free Quantification (LFQ) intensities of ME1 and ME2 in Col-0 and mutant lines grown under infected and mock condition. Statics between the abundances of ME1 and ME2 in (A) and between Col-0 and the *me*-mutant lines in (B) was assessed by Welch’s t-test (p < 0.05, *; p < 0.01, **; p < 0.001, ***; p < 0.0001, ****).

Principal component analysis (PCA) revealed genotype– and treatment-dependent differences in proteome composition. Under mock conditions, wild type and *me1* clustered separately from *me2* and *me1/me2*, consistent with the pronounced basal proteomic alterations associated with ME2 deficiency observed previously. Following infection, the two single mutants displayed more similar proteomic profiles, occupying an intermediate position between wild type and the double mutant (Fig. 6A). Comparisons within each genotype revealed the most pronounced separation between mock-treated and infected samples in *me1* (Fig. 6B). These patterns indicate genotype-dependent proteomic responses to infection.

**Figure 6.**
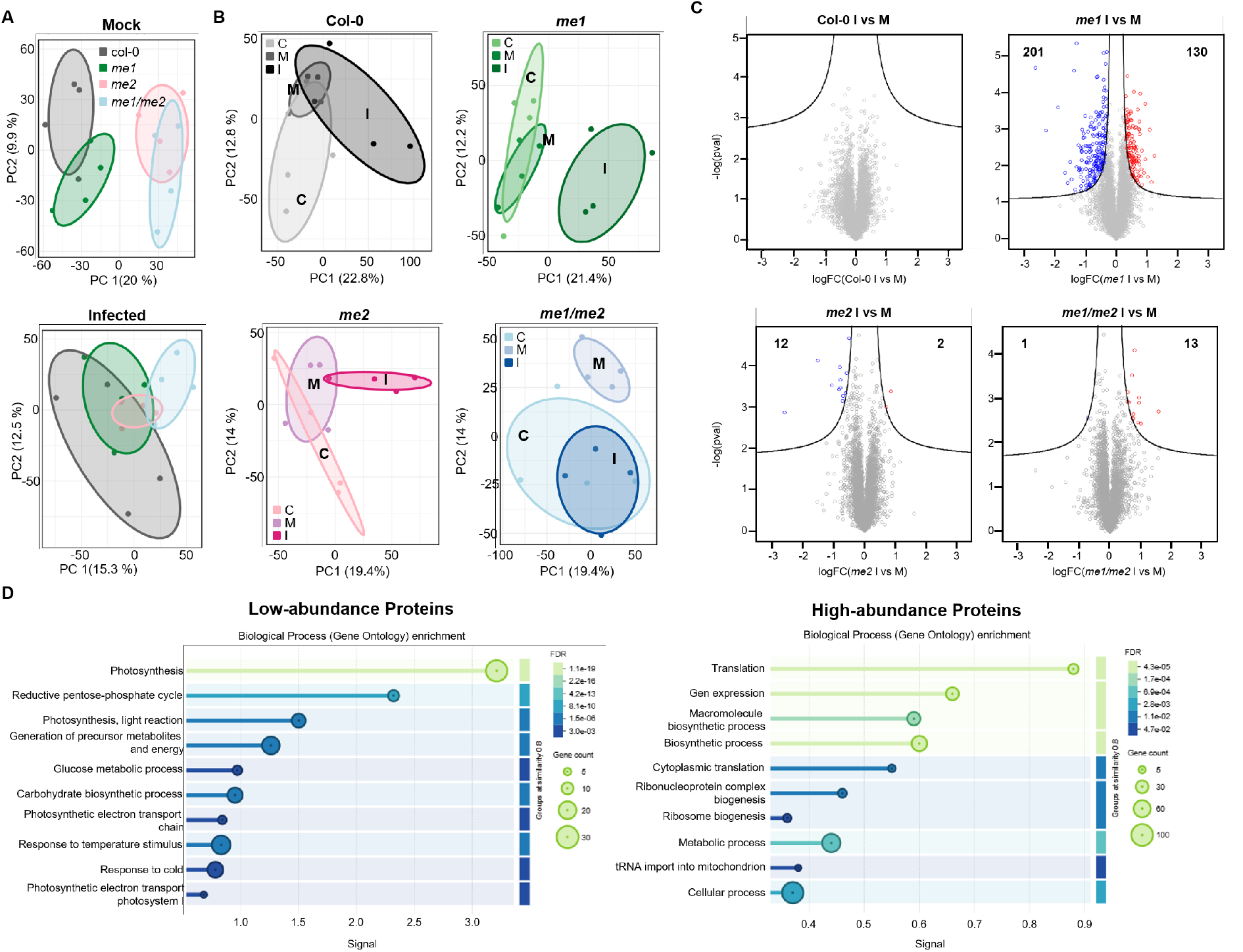
Proteomic analysis in *C. higginsianum* infected and mock-treated control leaves during the early stages of infection (2 dpi). **A**) PCAs of mock and infected samples of *me1*, *me2*, *me1/me2* and Col-0. Samples were harvested two days post infection, one hour before the light went off (+7) (n=4). **B)** PCAs from proteomic data under control (C), mock (M), and infected condition (I). **C)** After normalization, log_2_-transformed LFQ values of protein groups were used to assess differences in the cellular proteomes of *me1*, *me2* and *me1/me2* when compared against Col-0. Statistical testing employed an FDR ≤ 0.05 and S₀ = 0.1 using four independent biological replicates. Protein groups located in the areas above the black lines fulfil the significance threshold limits. The complete list of categories is provided in Suppl. Data 2. Red dots represent higher abundance protein groups and blue lower abundant. **D)** GO enrichment by STRING (v12.5) using higher and lower abundant proteins in *me1* infected compared to *me1* mock-treated.

Quantitative proteomic analysis identified 331 protein groups with significantly altered abundance in infected *me1* plants compared with mock-treated controls, including 130 that increased and 201 that decreased in abundance. In contrast, only 14 protein groups are significantly affected in *me2* (two increased and 12 decreased) and 14 in *me1/me2* (13 increased and one decreased; Fig. 6C). Thus, the absence of ME1 is associated with extensive infection-induced proteomic rearrangements, whereas ME2-deficient plants exhibit a markedly attenuated response at the proteome level. Given the substantial alterations already present in the basal proteomes of *me2* and *me1/me2*, the limited additional changes following infection may partly reflect their pre-existing proteomic state.

Gene Ontology (GO) enrichment analysis of the infection-responsive proteins in *me1* revealed distinct functional patterns. Proteins with increased abundance are enriched in processes associated with translation and gene expression, whereas those with decreased abundance are predominantly associated with the Calvin-Benson cycle, photosynthetic light reactions, and energy metabolism (Fig. 6D). These changes suggest a coordinated reorganization of cellular functions during infection, characterized by reduced investment in photosynthetic processes and increased abundance of proteins involved in gene expression and protein synthesis.

To investigate whether the constitutive proteomic alterations in ME2-deficient plants resemble the infection-induced response observed in *me1*, we compared proteins differentially abundant between infected and mock-treated *me1* plants with those differing between untreated *me2* or *me1/me2* plants and wild type under the same growth conditions. Seventeen proteins that decreased following infection in *me1* are also constitutively less abundant in both ME2-deficient genotypes (Table 1; Suppl. Fig. 4A; Suppl. Data 3). Among these, six are associated with photosynthesis, including Rubisco activase (RCA), CP12-1, and CP12-2 (Table1; Suppl. Fig. 4A and C; Suppl. Data 3). Conversely, four proteins show increased abundance in all comparisons, including choline/ethanolamine phosphotransferase 2, involved in phospholipid biosynthesis; ATP synthase E chain; and a chitinase family protein. However, no significantly enriched GO categories were identified using STRING (v12.5; Szklarczyk et al., 2025) (Suppl. Fig. 4B; Suppl. Data 3). These findings indicate that ME2 deficiency establishes a constitutive proteomic state that partially resembles the infection-induced response of *me1*, particularly with respect to reduced abundance of photosynthesis-associated proteins. However, all three NAD-ME mutants exhibit reduced fungal penetration, despite the substantially different extent of their basal proteomic alterations and infection-induced responses. The greater similarity of their proteomic profiles following infection suggests that shared infection-associated changes, rather than constitutive proteomic alterations, may contribute to the reduced fungal susceptibility observed across all three mutant genotypes.

**Table 1.**
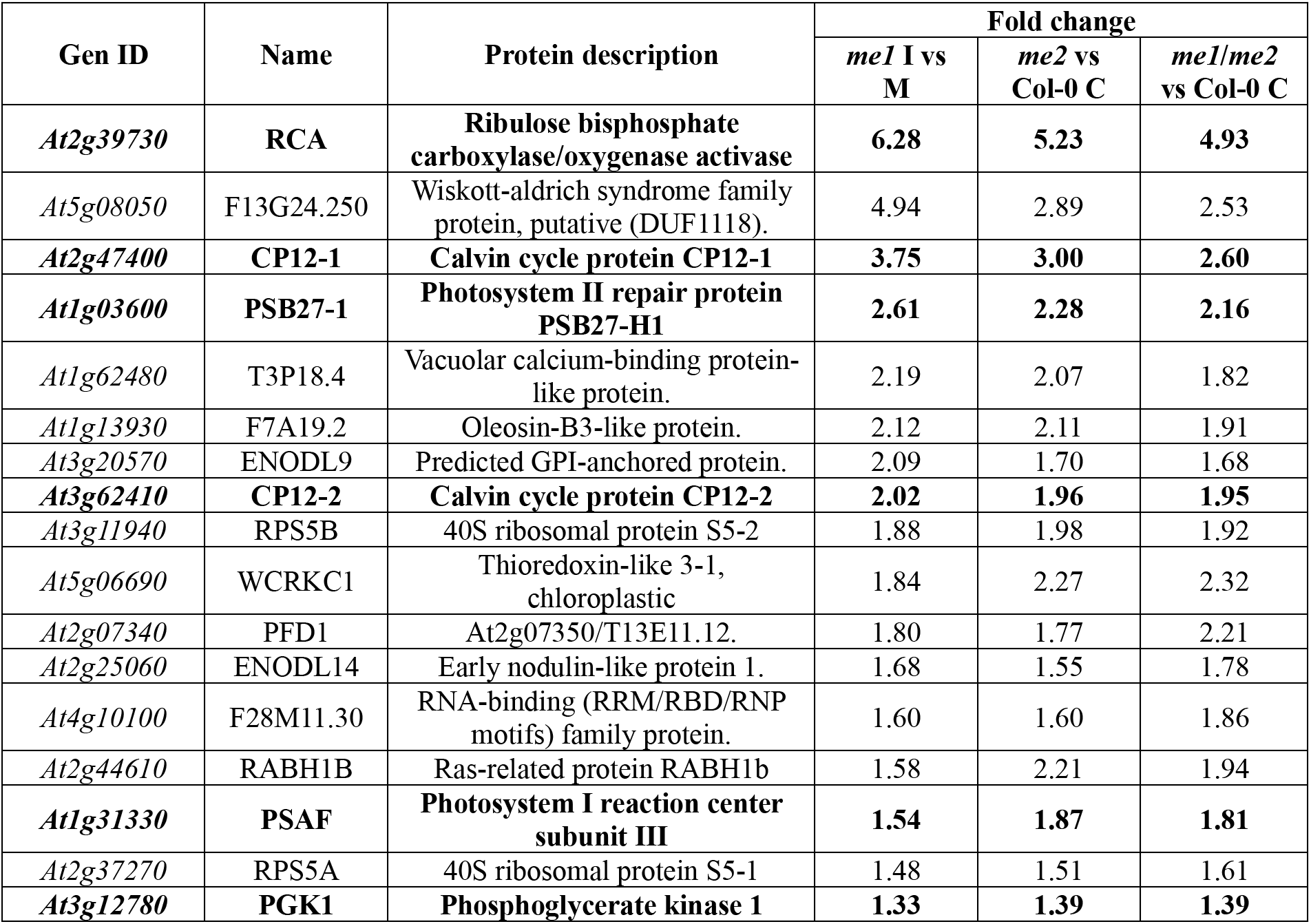
Proteins showing significantly lower abundance in common among infected me1 and me2 mutant lines under control conditions. The table lists proteins with reduced abundance shared between infected *me1* (I) compared with mock-treated *me1* (M), and *me2* mutant lines (*me2* and *me1/me2*) compared with Col-0. Fold changes indicate the magnitude of the decrease in protein abundance. In bold are marked the six proteins involved in photosynthesis.

| Gen ID | Name | Protein description | Fold change |  |  |
|---|---|---|---|---|---|
|  |  |  | *me1* I vs M | *me2* vs Col-0 C | *me1*/*me2* vs Col-0 C |
| ***At2g39730*** | **RCA** | **Ribulose bisphosphate carboxylase/oxygenase activase** | **6.28** | **5.23** | **4.93** |
| *At5g08050* | F13G24.250 | Wiskott-aldrich syndrome family protein, putative (DUF1118). | 4.94 | 2.89 | 2.53 |
| ***At2g47400*** | **CP12-1** | **Calvin cycle protein CP12-1** | **3.75** | **3.00** | **2.60** |
| ***At1g03600*** | **PSB27-1** | **Photosystem II repair protein PSB27-H1** | **2.61** | **2.28** | **2.16** |
| *At1g62480* | T3P18.4 | Vacuolar calcium-binding protein-like protein. | 2.19 | 2.07 | 1.82 |
| *At1g13930* | F7A19.2 | Oleosin-B3-like protein. | 2.12 | 2.11 | 1.91 |
| *At3g20570* | ENODL9 | Predicted GPI-anchored protein. | 2.09 | 1.70 | 1.68 |
| ***At3g62410*** | **CP12-2** | **Calvin cycle protein CP12-2** | **2.02** | **1.96** | **1.95** |
| *At3g11940* | RPS5B | 40S ribosomal protein S5-2 | 1.88 | 1.98 | 1.92 |
| *At5g06690* | WCRKC1 | Thioredoxin-like 3-1, chloroplastic | 1.84 | 2.27 | 2.32 |
| *At2g07340* | PFD1 | At2g07350/T13E11.12. | 1.80 | 1.77 | 2.21 |
| *At2g25060* | ENODL14 | Early nodulin-like protein 1. | 1.68 | 1.55 | 1.78 |
| *At4g10100* | F28M11.30 | RNA-binding (RRM/RBD/RNP motifs) family protein. | 1.60 | 1.60 | 1.86 |
| *At2g44610* | RABH1B | Ras-related protein RABH1b | 1.58 | 2.21 | 1.94 |
| ***At1g31330*** | **PSAF** | **Photosystem I reaction center subunit III** | **1.54** | **1.87** | **1.81** |
| *At2g37270* | RPS5A | 40S ribosomal protein S5-1 | 1.48 | 1.51 | 1.61 |
| ***At3g12780*** | **PGK1** | **Phosphoglycerate kinase 1** | **1.33** | **1.39** | **1.39** |

To assess whether these proteomic differences are accompanied by changes in primary metabolism, we analyzed metabolite profiles in rosettes collected two days after infection or mock treatment at four diel time points (−1, +1, +7, and +9 h relative to dawn). PCA did not reveal consistent separation among genotypes or treatments based on their overall metabolite profiles (Suppl. Fig. 5A and B). Although individual metabolites may exhibit genotype– or infection-dependent differences, the global analysis did not identify a common metabolic signature associated with reduced fungal penetration across the three mutants. Thus, the shared reduction in pathogen susceptibility is not readily explained by changes in bulk primary metabolite pools, but rather by proteomic changes in enzymes that may influence metabolic functions, which are not covered by the current-metabolic-profiling approach.

## DISCUSSION

### ME2-dependent accumulation of ME1 is associated with asymmetric proteomic remodeling

One of the important finding of this study is the pronounced asymmetry in the accumulation of NAD-ME subunits. In wild-type plants, ME2 is more abundant than ME1, and its abundance remains at approximately 75% of wild-type levels in *me1*. In contrast, ME1 protein abundance decreased to approximately 20% of wild-type levels in *me2*, despite the continued expression of its corresponding transcript. These findings indicate that ME2 is required for normal ME1 protein accumulation, suggesting that the abundance of the two subunits is regulated through distinct mechanisms. Interestingly, Tronconi et al. (2008) previously demonstrated that both single mutants retain residual NAD-ME activity, whereas the *me1/me2* double mutant completely lacks detectable activity, indicating that both subunits can form catalytically active homodimers in vivo. However, the contribution of the two subunits to total NAD-ME activity is markedly unequal, with the *me2* mutant retaining particularly low residual activity (less than 10% of wild-type activity). Our proteomic results provide a potential explanation for this asymmetry, as the pronounced reduction in ME1 abundance in *me2* would further limit the formation of catalytically active ME1 homodimers. The dependence of ME1 accumulation on ME2 presence could reflect impaired protein stability, altered assembly of NAD-ME complexes, or changes in post-transcriptional regulation.

The asymmetric dependence of NAD-ME subunit accumulation is accompanied by striking differences in basal proteome composition. Whereas the *me1* proteome remained relatively similar to wild type in control conditions, which indicates that ME2 is sufficient in the absence of environmental challenges, loss of ME2 resulted in extensive proteomic changes, with *me2* and *me1/me2* displaying highly similar profiles. These observations suggest that ME2 plays a particularly important role in maintaining proteome homeostasis under the conditions examined. Thus, the persistence of ME2 in *me1* may preserve sufficient NAD-ME function to limit broader metabolic and proteomic disturbances, whereas the strong reduction in ME1 abundance in *me2* may further compromise NAD-ME function.

Interestingly, the extensive proteomic remodeling observed in ME2-deficient plants includes reduced abundance of several proteins associated with photosynthetic carbon assimilation, such as RCA, CP12-1, and CP12-2, as well as components of the photosynthetic machinery. This suggests that perturbation of NAD-ME accumulation or action may influence chloroplast-associated processes, highlighting the close metabolic coordination between mitochondria and chloroplasts. However, these changes are not accompanied by detectable alterations in plant growth or chlorophyll fluorescence parameters under the experimental conditions examined. This apparent discrepancy may reflect compensatory mechanisms that maintain photosynthetic function despite changes in protein abundance or, more likely, indicate that the observed reductions in photosynthesis-associated proteins are insufficient to compromise light harvesting, photosynthetic electron transport, or overall photosynthetic performance. Alternatively, the affected proteins may become functionally important under environmental conditions like attempted fungal penetration, which impose greater demands on photosynthetic or respiratory metabolism.

Our study shows that he metabolic consequences of altered NAD-ME subunit composition may extend beyond mitochondrial NADH production. By converting malate into pyruvate, mitochondrial NAD-ME contributes to the coordination of organic acid metabolism and the availability of carbon intermediates for mitochondrial and cellular metabolic processes (Le et al., 2021; Le et al., 2022). Changes in NAD-ME activity could therefore influence metabolic exchange between compartments and indirectly affect chloroplast protein abundance. Future analyses of mitochondrial NAD-ME complex assembly, protein turnover, and compartment-specific metabolic fluxes will be necessary to clarify the mechanisms underlying the distinct effects of ME1 and ME2 deficiency.

### Disruption of mitochondrial NAD-ME reduces fungal penetration in the light of distinct proteomic responses

A second major finding of this study is that disruption of either mitochondrial NAD-ME subunit reduces penetration by the hemibiotrophic fungus *C. higginsianum*. All three mutant lines (*me1*, *me2*, and *me1/me2*) exhibit reduced fungal penetration, suggesting that mitochondrial NAD-ME facilitates successful pathogen establishment. As NAD-ME catalyzes the oxidative decarboxylation of malate to pyruvate, generating mitochondrial NADH, disruption of either subunit may alter metabolic and redox processes relevant to early plant-pathogen interactions. These alterations could influence host metabolic conditions required for fungal colonization or affect early immune signaling. However, whether the reduced susceptibility reflects impairment of a shared metabolic function or distinct contributions of the two NAD-ME subunits remains to be determined.

Recent evidence further supports a role for mitochondrial NAD-ME in plant-pathogen interactions. In *N. benthamiana*, silencing of *NAD-ME1* increased Bamboo mosaic virus accumulation and affected cytosolic NADH/NAD⁺ balance and defense-related gene expression, although the latter effects were relatively modest (Hou et al., 2025). Interestingly, these observations contrast with our findings that disruption of either mitochondrial NAD-ME subunit reduces fungal penetration in *A. thaliana*. These apparently opposite effects suggest that the contribution of mitochondrial NAD-ME to pathogen susceptibility depends on the nature of the host-pathogen interaction. Differences in pathogen lifestyle, nutritional requirements, and host metabolic responses may determine whether alterations in mitochondrial malate metabolism favor or restrict pathogen establishment. Thus, mitochondrial NAD-ME may influence plant susceptibility through context-dependent metabolic and redox processes rather than acting as a general positive or negative regulator of defense.

Despite their similar susceptibility phenotypes, the three NAD-ME mutants exhibited markedly different basal proteomes and proteomic responses to infection. Interestingly, a subset of proteins whose abundance decreased following infection in *me1*, including several photosynthesis-associated proteins, is also less abundant in ME2-deficient plants under non-infected conditions. This partial overlap suggests that the absence of ME2 establishes a constitutive proteomic state that partly resembles the infection-induced response observed in *me1*. However, also *me1* exhibits reduced fungal penetration despite lacking the extensive basal proteomic remodeling characteristic of ME2-deficient plants. This suggests that the reduced susceptibility shared by all three mutants does not require extensive constitutive changes in proteome composition. Together, these observations suggest that mitochondrial NAD-ME may influence early fungal establishment through metabolic or signaling processes.

Metabolite profiling provided further evidence that the shared susceptibility phenotype cannot be readily attributed to apparent steady state metabolite pools. Although genotype– and time-dependent differences were detected in several organic acids, amino acids, and sugars, the overall metabolite profiles did not reveal a consistent pattern associated with reduced fungal penetration across the three mutants. Nevertheless, metabolite pool sizes do not necessarily reflect metabolic fluxes or compartment-specific redox changes. The majority of the carboxylic acid pool is located in the vacuole. Hence, only a comparatively minor fraction of the carboxylate pools can respond dynamically to changes in metabolic flux. In fact, metabolic flux analysis is required to assess the underlying changes claimed here. Given the role of mitochondrial NAD-ME in malate metabolism and NADH production, alterations in mitochondrial redox homeostasis and its communication with other cellular compartments remain plausible mechanisms linking NAD-ME deficiency to pathogen susceptibility. Moreover, changes in mitochondrial metabolism could influence early defense-related processes, including reactive oxygen species production and downstream signaling pathways. Investigating these possibilities through compartment-specific redox measurements and early immune-response assays will be important for understanding how mitochondrial NAD-ME contributes to plant-pathogen interactions.

## MATERIAL AND METHODS

### T-DNA insertion lines

The plant lines used in this work include *A. thaliana* L. Heynh. ecotype Columbia (Col-0, wild type), along with homozygous T-DNA insertion lines Sail-374-A02 (*me1*) and Sail-291-C05 (*me2*), previously isolated by Tronconi et al. (2008) (Suppl. Table 1). The T-DNA insertion sites were verified by sequencing using specific primers (Suppl. Table 2). Double homozygous mutant was generated by genetic crosses. Homozygosity was confirmed by PCR using gene-specific and T-DNA left border primers (Suppl. Table 2).

### Plant growth

For all the experiments Col-0 and mutant lines were grown under at 22/19 °C (day/night) under short-day (SD: 8 h light/16 h dark) photoperiods, with light intensities of 60 μmol⋅m⁻²⋅s⁻¹ (LL). Plants for qRT-PCR analysis and metabolomics were grown in soil (Floradur ® Substrate Multiplication Seed B, Rüttger Abstohs Gärtner-Einkauf GmbH&Co.KG, Siegburg, Germany). Illumination was provided by a combination of Osram HO 80W/840 and HO 80W/830 bulbs. For proteome and metabolome analysis under infected, mock and control conditions, plants were grown in a 80:20 (v/v) mixture of soil (Frustorfer Erde Type P, Hawita, Germany) to expanded clay (Liadrain, Pautzfeld, Germany) in GroBanks (CLF, Wertingen, Germany) equipped with regular white light Gro-LED panels in the same light regime as described before. Plants were fertilized with 20 ml of 0.2% WUXAL super (Aglukon, Düsseldorf, Germany) one week prior to spray infection with *Colletotrichum higginsianum* or mock treatment as described below. Leaf material for proteome and metabolome analysis was harvested from 5 weeks old plants at 2 days post treatment. In all the cases, seeds were stratified at 4 °C for 3 days prior to transfer to growth conditions.

### RNA extraction and cDNA synthesis

Total RNA was extracted from rosette leaves of plants grown in SD under LL harvested 40 DAS. The harvest points were one hour before the light was on (−1), one hour after the light was on (+1) and one hour before the light was off (+7). Using the Universal RNA Purification Kit (Roboklon GmbH, Germany) according to the manufacturer’s instructions and eluted in 30 µl of RNase-free distilled water. RNA concentration and purity were determined using a NanoDrop Nabi™ UV/Vis spectrophotometer (MicroDigital Co. Ltd, South Korea), and RNA integrity was verified by agarose gel electrophoresis. Samples were stored at –20 °C until further use. To remove potential genomic DNA contamination, RNA samples were treated with the Ambion DNA-free™ DNA Removal Kit (Thermo Fisher Scientific, USA). Subsequently, 2 µg of RNA was reverse transcribed into complementary DNA (cDNA) using RevertAid Reverse Transcriptase (Thermo Scientific, USA) and an Oligo(dT) primers, following the manufacturer’s protocol.

### Reverse-transcription quantitative PCR

Reverse transcription quantitative PCR (RT-qPCR) was conducted using the KAPA SYBR FAST qPCR Kit (KAPA Biosystems Inc., Switzerland) to quantify the expression levels of *ME1* and *ME2* in cDNA samples derived from the different plant lines. Reactions were performed on a Mastercycler® ep realplex system (Eppendorf, Germany), and cycle threshold (Ct) values were obtained using the accompanying realplex software. Gene expression was normalized to the *ACTIN2* (*ACT2*) gene, which served as an internal reference.

### Photosynthetic parameters

Plants grown under SD at LL conditions 40 days after sowing were used for measuring photosynthetic parameters. PAM (Pulse Amplitude Modulation) measurements were performed using the IMAGING-PAM Maxi version (Walz, Germany) with a PAR (Photosynthetically Active Radiation) of 81 µmol m^-2^ s^-1^ following the method described by Martinez et al., (2026). Briefly, Plants had been dark-adapted for 20 min prior to each measurement. For each plant line, four biological replicates were analyzed, and five measurement points were selected from each replicate. An average of the 5 points from each biological replicate were used to calculate the values from each photosynthetic parameter. The maximum quantum yield efficiency of PSII (Fv/Fm = [Fm-F0]/Fm) was taken as a measure of PSII intactness, quantum Yield of PSII (ΦPSII) indicates the amount of solar energy converted to chemical energy to be used by the plant cell, the electron transport rate (ETR) shows the speed of the electrons in the electron transport chain and the non-photochemical quenching (NPQ).

### Metabolite analysis through GC-MS

To assess to the metabolomic profile in each line gas chromatography–mass spectrometry (GC-MS) was performed. All the lines were grown under LL in SD condition, and samples were taken from 40 days old plants, 1 hour before the light was on (−1), one hour after the light was on (+1) and one hour before the light was off (+7). The tissue was immediately frozen after harvesting, approximately 50 mg were used per each biological replicate. The frozen tissue was grinded using a TissueLyserII (QIAGEN, Germany) for 30 seconds two times at a frequency of 30 Hz. Soluble metabolites were extracted by a one phase extraction, using 1.5 ml a mixture of H_2_O:MeOH:CHCl3 (1:2.5:1) following the protocol by Fiehn et al. (2000). Samples were incubated 10 min at 4 °C shaking. After that, samples were centrifugated for 5 min at 4 °C at 20000 xg and 1 ml from the supernatant was transferred to a new tube. 30 µl from the samples were subsequently dried in a vacuum concentrator and derivatized for GC-MS analysis, as described by Gu (Gu et al., 2012). GC-MS measurements were carried out using a 5977B GC-MSD system (Agilent Technologies), following the procedure outlined by Shim et al., 2020 (DOI:10.3389/fpls.2019.01726). Metabolites were identified using MassHunter Qualitative Analysis software (version B.08.00, Agilent Technologies) by comparing acquired spectra with entries in the NIST14 Mass Spectral Library (https://www.nist.gov/srd/nist-standard-reference-database-1a-v14). A standard mixture containing all target compounds at a concentration of 5 µM was processed alongside the samples to serve as a quality control for response consistency and retention time referencing. Peak integration was performed using MassHunter Quantitative Analysis software (version B.08.00, Agilent Technologies). For relative quantification, metabolite peak areas were normalized to the fresh weight of the sample and to the peak area of the internal standard (Ribitol, Sigma-Aldrich). Statistics were performed for each time point using one-way ANOVA followed by Tukey’s post-hoc test.

### Plant infection and analysis

The *C. higginsianum* isolate MAFF 305635 (Ministry of Forestry and Fisheries, Japan) was cultivated on oatmeal agar plates previously described by Reckleben et al. (2025). Following a seven-day incubation period, the conidia were collected by rinsing the plates with 500 µl of deionized water. The conidia titer was subsequently adjusted to 2·10^6^ conidia · ml^-1^ using a hemocytometer, after which the suspension was used for spray infection on 35-days old plants at the end of the light phase, as previously described by Voll et al. (2012). High humidity was usually maintained until 2.5 days post infection. When fungal entry rates were determined under the microscope after staining fungal structures with trypan blue (Engelsdorf et al., 2017), or when samples for metabolome and proteome analysis were taken, high humidity was stopped at the time of harvest.

### Shotgun proteomic LC-MS analysis

Proteomic analysis was performed on plants grown under SD conditions at LL in control, mock and infected conditions. Whole rosettes were harvested 2 days post infection one hour before the end of the photoperiod. Infection was done following the same methodology explained previously. Four biological replicates with 3-5 mg of dry weigh per genotype were collected. Protein extraction and digestion was carried out using a single-pot, solid-phase-enhanced sample preparation (SP3) protocol (Hughes et al., 2019), which was adapted to meet challenges inherent to plant samples (Mikulášek et al., 2021) as outlined in (Martinez et al., 2026). Each extract was treated with 2 µg of pre-activated, sequencing grade Trypsin (V5111, Promega). Concentrations of extracted tryptic peptides were determined in a plate reader (Multiskan Sky, Thermo Fischer Scientific) using the Pierce Quantitative Colorimetric Peptide Assay Kit (Thermo Fisher Scientific, Dreieich, Germany) following the manufacturer’s instructions. Peptides were dried using a vacuum concentrator and resuspended in 0.1% (v/v) formic acid (FA) to yield a peptide concentration of 10 ng µl^-1^. Twenty microliters of the suspension were loaded on Evotips (Evosep) according to the manufacturer’s instructions for desalting and subsequent separation using an EvoSep One HPLC (Evosep). Peptides were dried using a vacuum concentrator and resuspended in 0.1% (v/v) formic acid (FA) to yield a peptide concentration of 10 ng µl^-1^. Twenty microliters of the suspension were loaded on Evotips (Evosep) according to the manufacturer’s instructions for desalting and were subsequently separated using an EvoSep One HPLC (Evosep). Peptides bound to the Evotip resin were eluted from a 15 cm Aurora Elite CSI column (inner diameter 75 µm, particle size 1.7 µm, pore size 120 Å, IonOpticks, Fitzroy, Australia) at a temperature of 50 °C using the Whisper Zoom 40 SPD method. Eluting peptides were analyzed in a timsTOF Pro mass spectrometer (Bruker, Germany) using the pre-installed ‘dia-PASEF_1sec_cycletime_short’ method.

Acquired spectra were queried against an in-house spectral library containing TAIR10 and ARAPORT11 database entries considering mRNA editing events using the DIA-NN software (Demichev et al., 2020) version 2.2.0. Default parameters were applied with the following exceptions: oxidation of methionine and acetylation of N-terminus were enabled as variable modifications, precursor charge was limited to 2-4, and the m/z range was set to a maximum of 1700. Mass accuracies were set to 15 ppm, the ‘match between runs’ function was enabled and the ‘double-pass’ neural network function was selected for machine learning. Protein abundance within each sample was estimated by computing intensity-based abundance quantification (iBAQ) values (Schwanhäusser et al., 2011) calculated by DIAgui application (Gerault et al., 2024) in R Studio version 4.5.0. The peptide length restriction was set to 7-30 for this.

The resulting ‘proteingroups.txt’ file was complemented with predicted subcellular protein localization using the SUBAcon algorithm (Hooper et al., 2014). The classified table was subsequently submitted to statistical analysis by means of the Perseus software (Tyanova et al., 2016) version 2.1.2.0 in the form of two-sample t tests (*p* < 0.05). For this, protein groups matching ‘only identified by site’, ‘potential contaminant’ and ‘reverse’ were removed before LFQ values were log_2_-transformed. Protein groups with less than three values in each sample group were removed and missing values were imputed from normal distribution using a width of 0.3 and a down-shift of 1.8. A width normalization was performed and volcano plots were produced using default parameters (FDR, 0.05; S_0_, 0.1).

Venn Diagrams were built using Venny2.1 (https://bioinfogp.cnb.csic.es/tools/venny/index.html) and GO term enrichment analysis was performed using STRING (v12.5, Szklarczyk et al., 2025).

### Author contributions

V.M., L.V. and M.P.M. conceived the study. M.P.M., A.F., I.N., N.D., J.S., C.R. and P.W. performed experiments and collected data. V.M., L.V., M.P.M., N.D. and H.E. analyzed the data and interpreted results. V.G.M and M.P.M. wrote the manuscript with input from L.V., N. D. and H.E.. V.M. and L.V. supervised the project.

### Data availability

All MS raw files acquired in this study, along with the corresponding MaxQuant search results can be downloaded from the MassIVE FTP server ftp://MSV000104072@massiveftp.ucsd.edu. Or on the MassIVE website using the MassIVE identifier: MSV000104072.

## Acknowledgements

This work was supported by the Deutsche Forschungsgemeinschaft (DFG, German Research Foundation) through the PAK918 initiative “Plant Mitochondria in New Light” (MA2379/14-3) to V.M., grant EU54/7-1 to H.E., and infrastructure grant INST 187/791-1 FUGG. We thank the CEPLAS Metabolism & Metabolomics Laboratory (CMML) at Heinrich Heine University Düsseldorf for technical assistance. CMML is supported by the DFG through the Cluster of Excellence CEPLAS (EXC-2048/1; Project ID 390686111).

