## Supplementary Figures for "Mitochondrial NAD-malic enzyme subunits modulate early fungal penetration and differentially shape the *Arabidopsis* proteome"

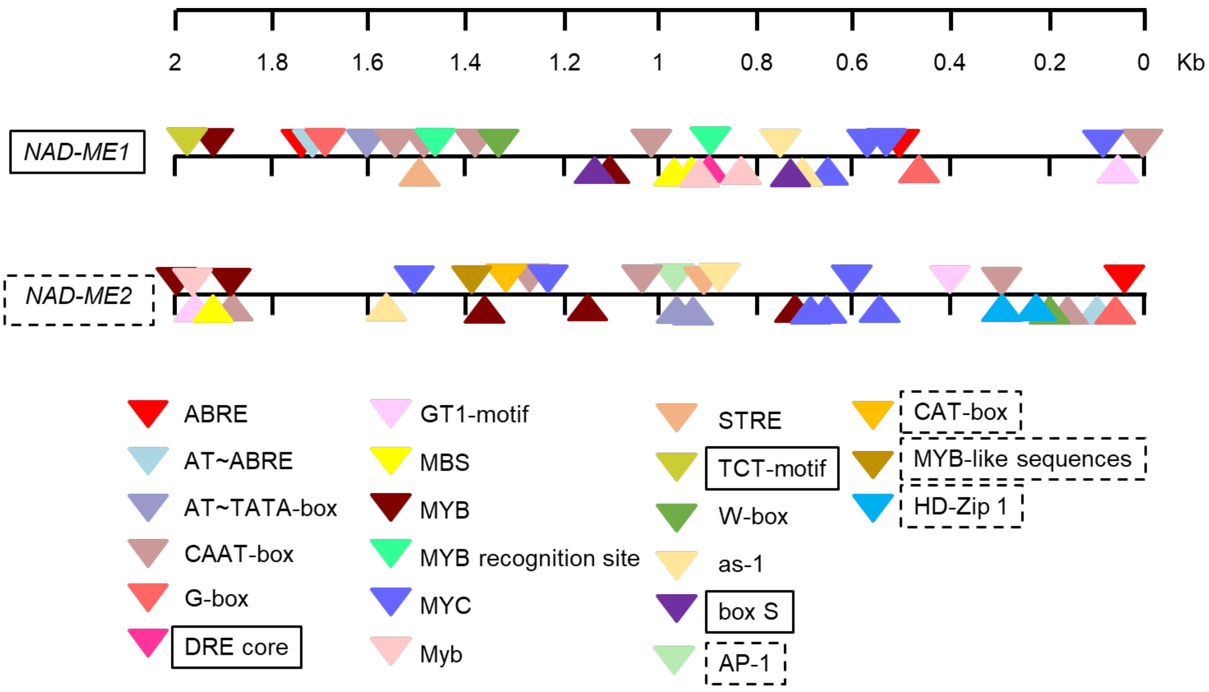


**Supplementary Figure 1. In silico analysis of the *ME1* and *ME2* promoter regions.** Schematic representation of the cis-regulatory elements identified using PlantCARE in the 2-kb regions upstream of the ATG start codon of *ME1* and *ME2*. Elements uniquely identified in either the *ME1* or *ME2* promoter region are highlighted with boxes.

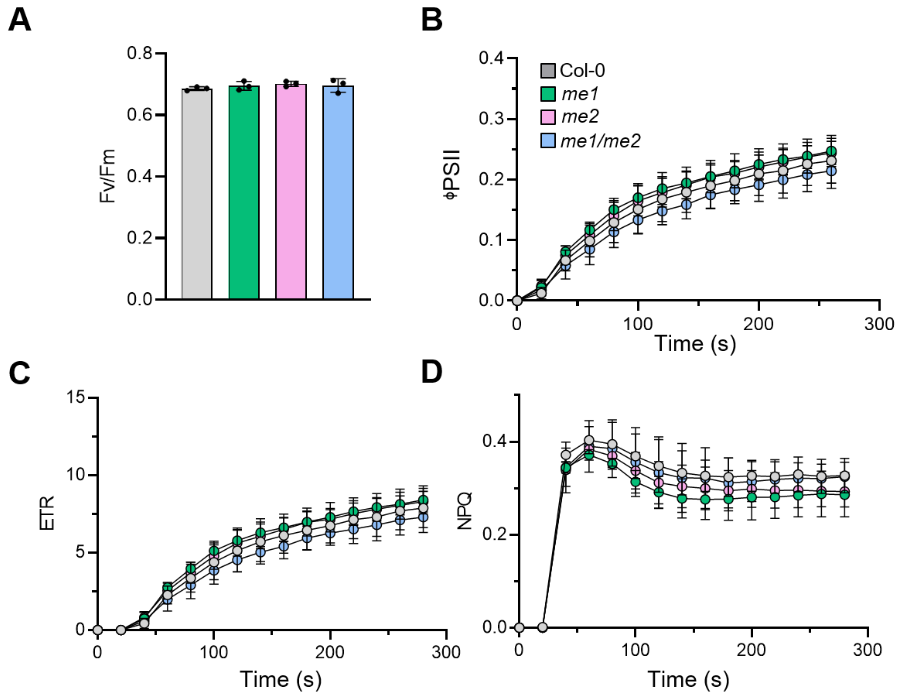


**Supplementary Figure 2**. **Photosynthetic performance of plants grown in LL-SD**. Photosynthetic parameters derived from chlorophyll fluorescence, **(A)** Fv/Fm (maximum quantum yield of PSII), **(B)** ΦPSII (operational quantum yield of PSII), ETR (electron transport rate), and **(C)** NPQ (non-photochemical quenching), were measured in 5-week-old plants grown at LL. Error bars represent ± SE (n=4). Four biological replicates were used per genotype, with five randomly selected spots from the middle of the leaf blade. No statistical difference was detected between the genotypes.

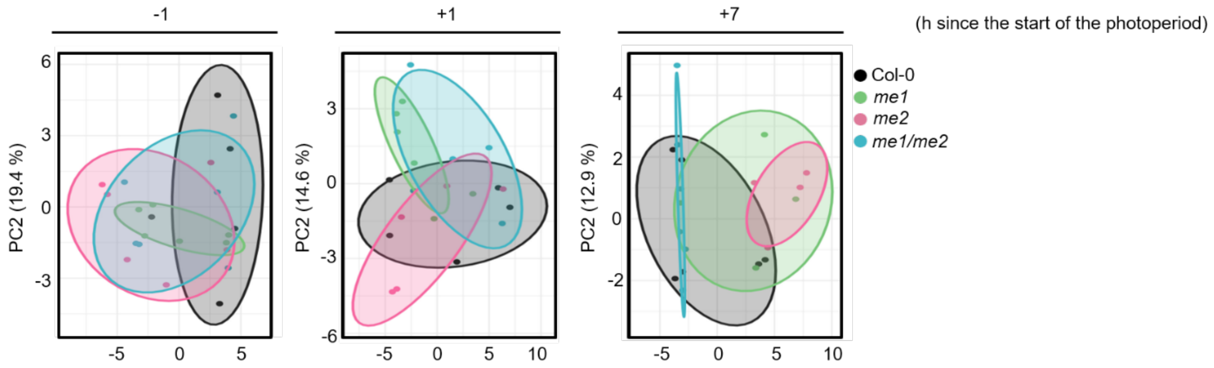


**Supplementary Figure 3. Principal component analysis (PCAs) from metabolomic data of *me* mutant lines and Col-0 harvested at different time points.** PCA analysis of metabolomic data from mutant lines and Col-0. Whole rosettes were harvested one hour before lights on (-1), one hour after (+1) and one hour before the photoperiod ends (+7) (n=4-5).

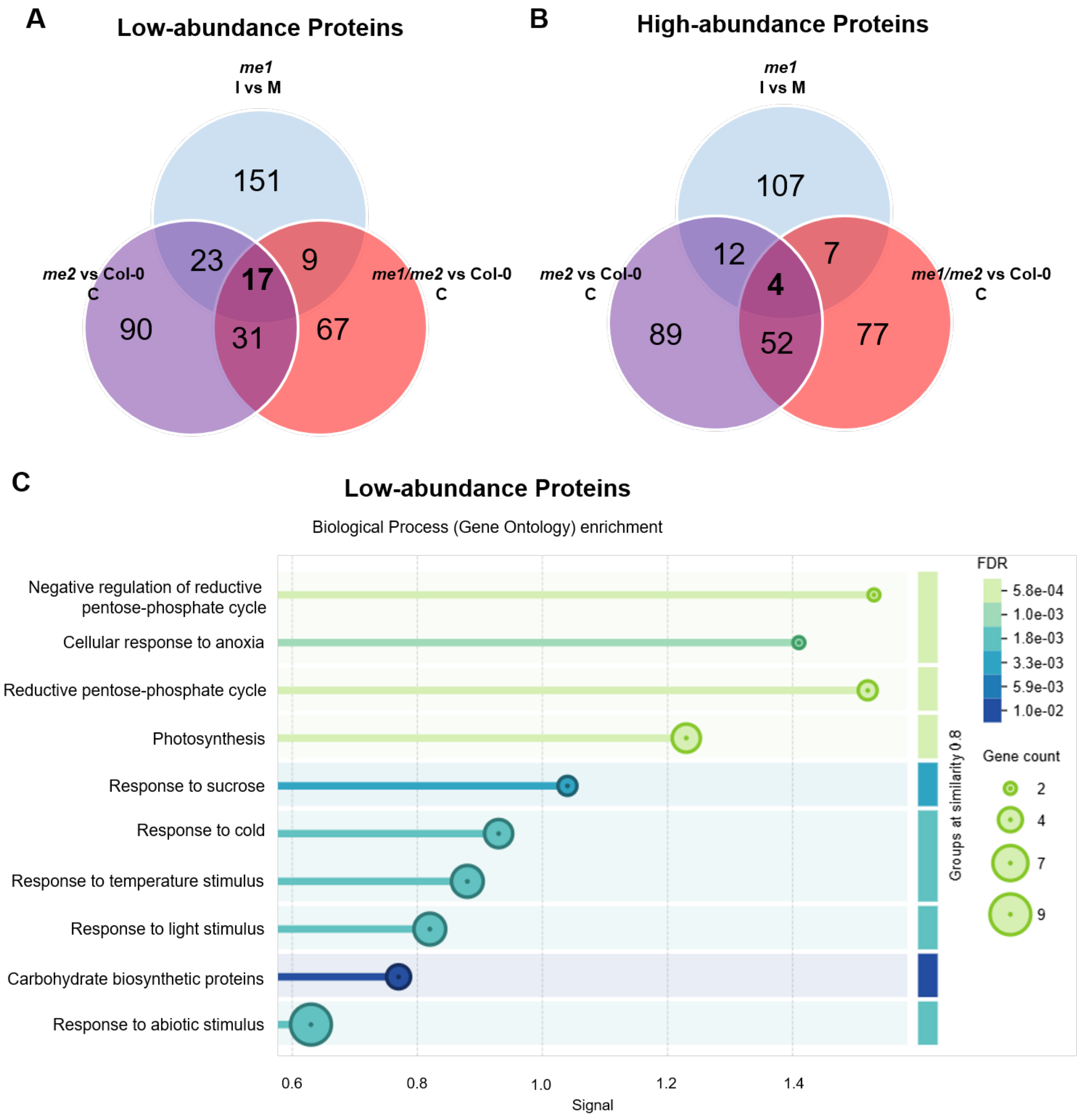


**Supplementary Figure 4. Comparison in proteomic changes in *me1* infected and *me2* mutants against col-0 under control conditions.** Venn Diagrams indicating unique and common **(A)** lower and **(B)** higher abundance protein groups in *me2* lines against Col-0 under control condition (C) and in *me1* infected (I) against mock (M). **C)** Gene Ontology enrichment using STRING (v12.5) with the 17 common low abundance proteins between infected *me1*, and *me2* and *me1/me2* against Col-0 under control condition selected in **A**.

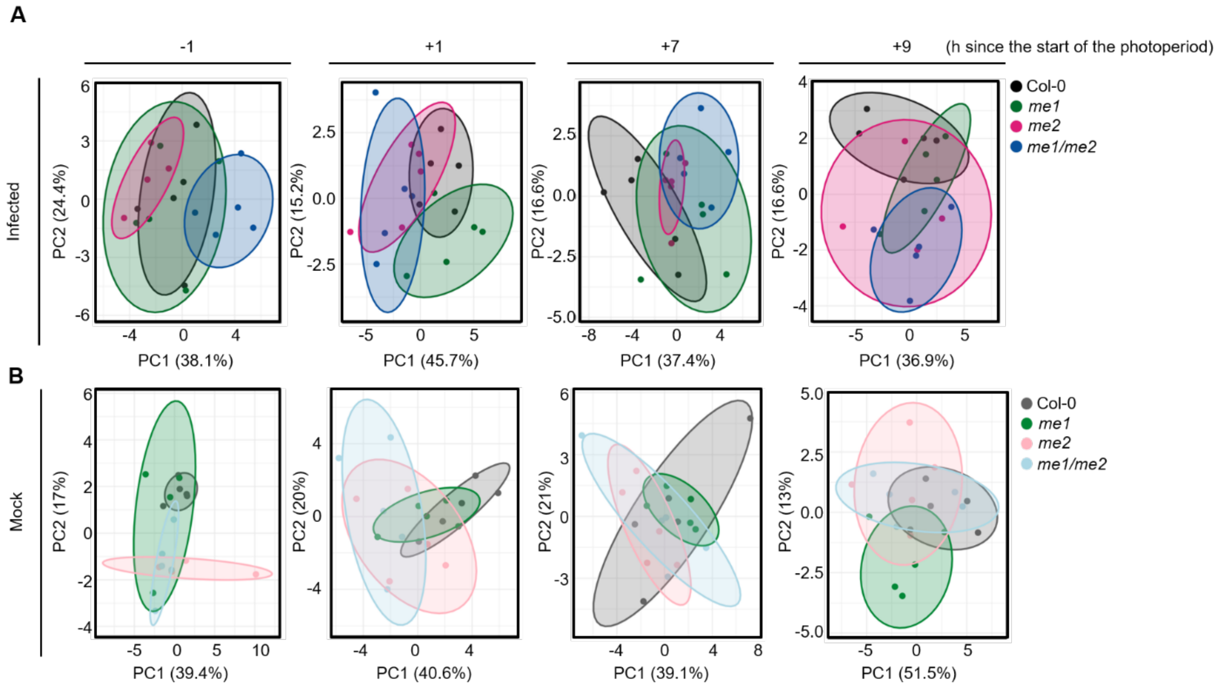


**Supplementary Figure 5. Principal component analysis (PCAs) from metabolomic data of *me* mutant lines and Col-0 harvested at different time points.** PCA analysis of metabolomic data from mutant lines and Col-0 **(A)** 2 days after fungal infection (infected), and **(B)** in mock conditions with the same high humidity (mock). Fully expanded leaves were harvested one hour before lights on (-1), one hour after (+1), one hour before the photoperiod ends (+7) and one hour into the night period (+9) (n=4-5).
