## Supplementary Tables for "Mitochondrial NAD-malic enzyme subunits modulate early fungal penetration and differentially shape the *Arabidopsis* proteome"

**Supplementary Table 1.** Information on the T-DNA insertion lines used in the work. s. a. = see above; ko = knock out.

| Mutant (Type) | Gene | Gene locus | Insertion line | Reference |
|---|---|---|---|---|
| *me1* (ko) | *ME1* | *At2g13560* | Sail-374-A02 | Tronconi et al., 2008 |
| *me2.1* (ko) | *ME2* | *At4g00570* | Sail-291-C05 | |
| *me1*/*me2* | s. a. | s. a. | s. a. | |

**Supplementary Table 2.** List of primers used in this study.

| **Name** | **Sequence (5'- 3')** | **Gene/T-DNA Insertion** | **Use** |
|---|---|---|---|
| me-1-Fw-1 | ACGATGACGGAGAGAAATCGT | *ME1* | PCR |
| me-1-Rv-1 | CTGGACATCATCATTGAACAT | | |
| me-2-Fw-1 | GACCTGTGTACAGCAATGTGATCG | *ME2* | |
| me-2-Rv-1 | CCTGGACATCATCATTGAACATGC | | |
| act2-Fw-1 | TAACTCTCCCGCTATGTATGTCGC | *ACT2* | |
| act2-Rv-1 | GAAGCAAGAATGGAACCACCG | | |
| Sail | TAGCATCTGAATTTCATAACCAATCTCGATACAC | *ME1 / ME2* | |
| me-1-Fw-2 | AAGATGAAGATCGTTGTTGCTG | *ME1* | qRTPCR |
| me-1-Rv-2 | CATCAACCACCCAAAATTGACT | | |
| me-2-Fw-2 | TTGGTGTAACTAAGATGGCAGT | *ME2* | |
| me-2-Rv-2 | CGTATCTCAGCTGGATTTTTGG | | |
| act2-Fw-2 | CTTGCACCAAGCAGCATGAA | *ACT2* | |
| act2-Rv-2 | CCGATCCAGACACTGTACTTCCTT | | |

**Supplementary Table 3. In silico analysis of the *ME1* and *ME2* promoter regions.** Schematic representation of the cis-regulatory elements identified using PlantCARE in the 2-kb regions upstream of the ATG start codon of *ME1* and *ME2*.

| cis- Elements | Position upstream of the transcript start site (bp) | Strand | Sequence | Gene | Function |
|---|---|---|---|---|---|
| ABRE | -404 | + | ACGTG | *ME1* | abscisic acid responsiveness |
| | -737 | - | AACCCGG | | |
| | -1644 | + | ACGTG | | |
| | -405 | - | CACGTG | | |
| | -70 | - | ACGTG | *ME2* | |
| AT-ABRE | -1645 | + | TACGTGTC | *ME1* | |
| | -72 | - | TACGTGTC | *ME2* | |
| AP-1 | -947 | + | TGAGTTAG | *ME2* | heat stress and heavy-metal defense |
| CAT-box | -1261 | + | GCCACT | *ME2* | meristem expression |
| G-box | -70 | - | TACGTG | *ME2* | light responsiveness |
| | -1645 | + | TACGTG | *ME1* | |
| | -405 | - | CACGTG | | |
| GT1-motif | -47 | - | GGTTAA | *ME1* | light responsiveness |
| | -1954 | - | | *ME2* | |
| | -380 | + | | | |
| DRE core | -628 | - | GCCGAC | *ME1* | dehydration |
| HD-Zip 1 | -215 | - | CAAT(A/T)ATTG | *ME2* | differentiation of the palisade mesophyll cell |
| | -163 | - | | | |
| MBS | -870 | - | CAACTG | *ME1* | drought-inducibility |
| | -792 | - | | *ME1* | |
| | -1970 | - | | *ME2* | |
| MYB | -1936 | + | CAACCA | *ME1* | growth, development, and stress responses |
| | -1042 | - | | | |
| | -1996 | + | CAACCA | *ME2* | |
| | -622 | - | | | |
| | -1339 | - | TAACCA | | |

| | | | | | |
|---|---|---|---|---|---|
| | -1890 | - | CAACAG | | |
| | -1143 | - | | | |
| MYB recognition site | -1410 | + | CCGTTG | *ME1* | secondary metabolism and abiotic stress |
| | -813 | + | | | |
| MYB like sequence | -1339 | - | TAACCA | *ME2* | |
| MYC | -623 | - | CATTTG | *ME1* | hormones signalling |
| | -489 | + | | | |
| | -581 | + | | | |
| | -11 | + | | | |
| | -1504 | + | | *ME2* | |
| | -581 | + | | | |
| | -635 | - | | | |
| | -545 | - | | | |
| | -1272 | + | | | |
| | -626 | - | | | |
| Myb | -870 | - | CAACTG | *ME1* | |
| | -792 | - | | | |
| | -1970 | - | | *ME2* | |
| STRE | -1302 | - | AGGGG | *ME1* | stress response |
| | -886 | + | | *ME2* | |
| TCT-motif | -1922 | + | TCTTAC | *ME1* | light responsiveness |
| W-box | -1396 | - | TTGACC | *ME1* | binding site for WRKY |
| | -136 | + | | *ME2* | |
| as-1 | -727 | + | TGACG | *ME1* | response to salicylic acid auxins, and oxidative stress |
| | -696 | - | | | |
| | -1575 | - | | *ME2* | |
| | -862 | + | | | |
| Box S | -1129 | - | AGCCACC | *ME1* | response to fungal elicitors and biotic stress |
| | -687 | - | | | |
